# Integration of external biomass reactions into existing metabolic models

**DOI:** 10.1101/2022.08.01.502323

**Authors:** María Moscardó García, Maria Pacheco, Thomas Sauter

## Abstract

Currently, seven biomass objective functions have been defined in human metabolic reconstructions. The integration of published biomass reactions into alternative models can contribute to the prediction power of the model. Thus, in this work, we present a workflow to integrate reactions and biomass functions originating from several genome-scale reconstructions into models other than their home models. Additionally, a benchmark to identify the biomass that confers the highest prediction accuracy in terms of gene essentiality and growth predictions is provided.

**For complete details on the use and execution of this protocol, please refer to Moscardó García et al. (2021)**.

## Before you begin

Computational approaches such as context-specific metabolic models reconstructed from patients or cell line RNA-seq data are becoming more and more popular to guide and complement biological experiments. Nevertheless, one main issue, which directly impacts the quality of the context-specific models and thus their applications, remains, the formulation of the objective function. To establish an objective function, one should answer the questions: What are the main tasks of the tissue, cell, or organism, and what are the metabolites or macromolecules that must be produced to fulfil these tasks? While for cancer and high-proliferative cells, growth via the production of biomass might be a suitable objective, for cells that have slow proliferation speeds or for tissues in a pluricellular organism other functions must be considered such as energy production, the production and absorption of key metabolites, or minimisation of internal fluxes. Considering biomass production, there is no agreement on what quantity in mM of amino acids, nucleotides, sugars, and fatty acids would be required to produce 1 g of biomass. Several formulations can be found within GEMs from the same organism with different required mass-fractions of each metabolite (how much mass weight of the biomass metabolites in g/mmol are required to produce 1 g of biomass but also regarding the biomass composition). Some metabolites are omitted in some formulations. Hence, in the present study that is based on the paper: Importance of the biomass formulation for cancer metabolic modelling and drug prediction by Moscardo Garcia et al, published in 2021 in iScience, we provide a protocol to add an objective function from a genome-scale reconstruction (GEM) into a different GEM or model and to test the predictive power of the GEM with this new reaction against experimental data. The pipeline can be applied to any organism of interest by using the corresponding GEMs and experimental data for the generation and validation of the results, respectively. However, in the present protocol, we explain first how to integrate human biomass functions into the input model of interest. Then how to build a cancer model using rFASTCORMICS and transcriptomic data from the CCLE repository. And, finally, how to test the predictive power of the cancer model in terms of essentiality against high throughput CRISPR-Cas 9 data and predicted growth using experimentally obtained doubling.

However, this protocol is not restricted to the biomass reaction and could be followed to integrate any objective function or reaction of interest in any GEM. Technical details on transcriptomics data acquisition can be found in Barretina et al., 2012. Similarly, the experimental gene essentiality and growth rate data protocols can be found in DepMap Achilles 19Q1, 2019 and O’Connor et al., 1997, respectively. The software setup described here was deployed on Windows 10 distribution. However, all referred tools and packages function properly across different platforms. Detailed instructions on the installation process for macOS and Windows can be found on https://github.com/sysbiolux/rFASTCORMICS.

## Installation of analysis tools

### Timing: 1 h

1. Install MATLAB on your computer system. MATLAB is a commercial software and a programming environment that can be downloaded from https://nl.mathworks.com/. Student licenses are available.
  a. Install the following toolboxes during the MATLAB installation or as AddOns later:
    I. Statistics and Machine learning Toolbox.
    II. Curve Fitting Toolbox.
2. Install the IBM solver CPLEX. This solver is free for academics upon registration on https://www.ibm.com/academic/topic/data-science. MATLAB should be launched and CPLEX should be added to the path via the command add files(). To verify that the installation was done properly type *Cplex()* in the Command window. Once the CPLEX solver has been installed, its selection is required using:

~~~
> changeCobraSolver(‘ibm_cplex’)
~~~ **CRITICAL:** CPLEX has an installer that will install the solver in the program files. Once the installation is done check if a MATLAB folder is present in the path C:\Program Files\IBM\ILOG\CPLEX_Studio201\cplex\MATLAB. If the MATLAB file is missing CPLEX has to be reinstalled and another version might need to be used.
3. Download rFASTCORMICS. This pipeline requires a context-specific building algorithm called rFASTCORMICS (Pacheco et al., 2019) which builds context-specific models (tissue-specific models and condition-specific models such as a cell in hypoxia or cancer cells) from genome-scale reconstructions and RNA sequencing data. The guiding principle behind the FASTCORE family is compactness of the reconstructed networks. The algorithm builds a consistent network around a set of core reactions, which are supported by the gene expression data. If a pathway or a reaction is not supported by the data or it has a too low expression value, it will not be included. Thus, in the discretisation step, two thresholds are determined based on the data distribution for each sample by fitting 2 Gaussian curves to the log-transformed data. The top of the blue Gaussian curve in Figure 1 corresponds to the expression threshold for this sample. The top of the red Gaussian corresponds to the not-expression threshold. Genes with expression values above the expression threshold are expressed (core genes), whereas genes with expression below the not- expression threshold are not expressed. Reactions that are under the control of core genes constitute the core reactions set, except in the case where the reactions are under the control of complexes of proteins and one or more genes coding a subunit of this complex is not expressed. If all the genes that control reactions are determined to be not expressed by the discretisation step, the reaction (inactive reaction) is removed from the model along with all reactions, including core reactions, that are no longer consistent because of the removal. A version of FASTCORE is then run on the remaining network to build a compact consistent network that contains all the reactions that did not depend on the presence of an inactive reaction.

**Figure 1.**
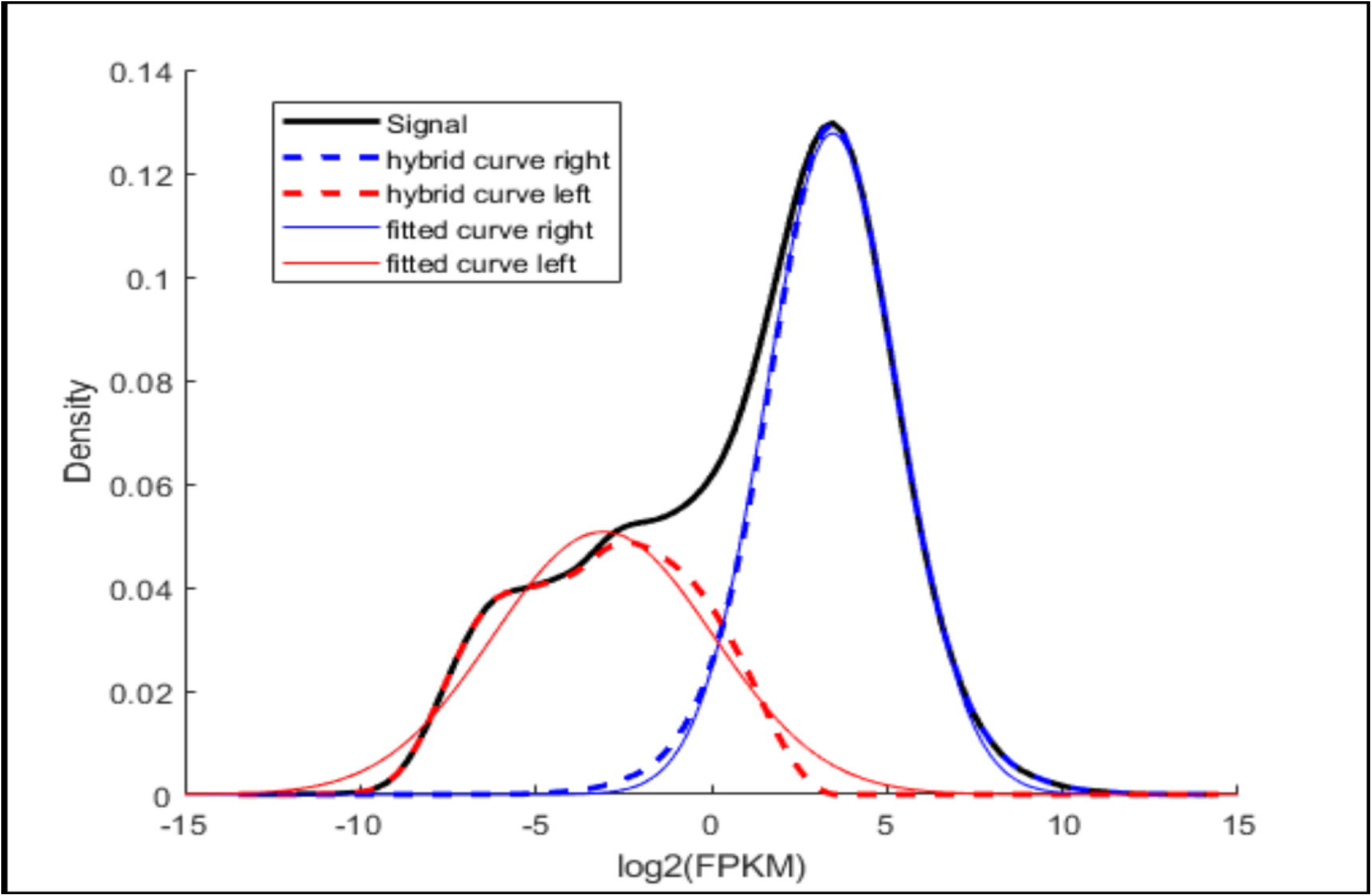
Quality control plots generated by rFASTCORMICS. These plots allow checking the distribution of the data for each gene expression sample. First, we have the fitting of the Gaussian curves (left) for the log2FPMK values. Afterwards, a z-score is obtained and genes are classified as expressed or not expressed (right, green lines). rFASTCORMICS can be retrieved from https://github.com/sysbiolux/rFASTCORMICS and, after downloading it, it should be set in the MATLAB path with the command addpath (). rFASTCORMICS creates quality control plots that show the distribution of each sample (gene expression data) and where the gene expression threshold was set for each sample. Ideally, the plot should look like the following example (Figure 1):
4. Download COBRA toolbox. The COBRA toolbox is required to assess the performance of each model with the tested biomass reaction. It can be directly retrieved from https://opencobra.github.io/cobratoolbox/stable following the installation guide. It prior requires the installation of git from https://git-scm.com/download/win. Once the installation is done, MATLAB is launched and the toolbox can then be initialised by running the command *initCobraToolbox* in the command window.

## Download required data

### Timing: 30 min

5. Download input data. Gene expression data can be downloaded and integrated into the input model to obtain context-specific models using rFASTCORMICS. In this particular case, RNA sequencing data from several cancer cell lines was retrieved from https://depmap.org/portal/download/. However, any other RNA sequencing dataset measured in Fragments Per Kilobase of transcript per Million mapped reads (FPKM) value could be used following the same pipeline. The data should be provided as a matrix with the rows corresponding to the gene identifiers or gene symbol and the columns to the different conditions and should contain only the FPKM values. Another variable named rownames in our code is obtained from the rows of the expression matrix, so it contains the gene identifiers or gene symbols present in the dataset. The “rownames” variable should be provided as a cell array and should have the same number of rows as the matrix. The “columnnames” should be provided as a cell array that contains the names of the conditions or the names of the samples and should have the same length as the columns in the matrix. **Note**: This example was run with rFASTCORMICS since RNA sequencing data was used as input. However, microarray data could also be used as input data using FASTCORMICS (Pacheco et al., 2015) as the context-building algorithm instead.
6. Download input model. There are several repositories where models are stored. The most common repositories are the BiGG database (http://bigg.ucsd.edu/), the VMH website (https://www.vmh.life/), and the Metabolic Atlas https://metabolicatlas.org/. Alternatively, the models can be retrieved from the supplementary files of published papers or the GitHub repositories of the authors. Overall, the recommended format to obtain the models is SBML, which can be read on MATLAB using the COBRA toolbox function readCbModel.m. **Note:** In the present example Recon 2 (Thiele et al., 2013) was used as input reconstruction. But other generic reconstructions could be used as input such as HMR 2.0 (Mardinoglu et al., 2014) or Human 1 (Robinson et al., 2020), as well as GEMs for other organisms such as plants. It is generally advisable to use to use more recent versions of reconstruction as issues present in earlier version might have been corrected in more recent reconstructions. Although, for some applications and in some studies, older more compact models yield slightly better results.
7. Download validation data. Validation data will be used to compare the model predictions against experimental values. In this study, two possibilities to assess the model performance are suggested: essential gene and growth rate predictions. Usually, CRISPR-Cas 9 datasets are used for the validation of essential gene predictions. Thus, CRISPR-Cas 9 data for the samples and genes considered in the gene expression dataset and obtained under the same experimental conditions as those mimics during the gene essentiality assay is required. On the other hand, experimental growth rates should be obtained for the samples under study (same cell lines or mutants considered in the gene expression data) and with the culture conditions mimic during the simulation analysis. Additionally, the growth rate analysis requires exometabolomic data, to constrain the model used for simulation and improve the predictions. Such data will be obtained from experimental analysis tailored to our simulated system (in terms of culture conditions) and the defined bounds should be tailored to the input model. Most biomass are given in h-1. In principle, by multiplying the simulated growth (without the default bounds) value by the coefficients mM gDW-1, one can calculate the simulated uptakes rates for the metabolites present in the biomass formulation. The simulated uptake rates are compared to the measured ones and the bounds of the uptake metabolites are adjusted until the simulated uptake rates correspond to the measured rates. In practice, the process is more complex as experimental measurements are not always in the correct unit, notably the experimental data is often expressed in mM per cells, due to shortcomings in some metabolic models the fixing of the bounds for some metabolites might result in no solution. Hence in some cases, some constraints need to be relaxed. Since the definition of such bounds is out of the scope of this paper, we refer the reader to previous publications explaining how to define these bounds (Jain et al., 2012, Zielinski et al,2017). Particularly, CRISPR-Cas 9 data (DepMap Achilles 19Q1, 2019) and experimentally measured growth rates (O’Connor et al., 1997) were used for validation in this study. For other organisms or applications for which exometabolite data might not be available, flux rates can be obtained from the literature. Another option is to predict the growth on different carbon sources or minimal media and to compare the results with experimentally observed values and in different conditions such as in hypoxia versus normoxia. Concerning essential genes, experimental settings that compare the capacity of mutants to survive or to survive on a given media can be simulated and compared to experimental data.

## Step-by-step method details

### Step 1: Extract the reactions of interest

#### Timing: 5 min

The goal of the study is to collect currently published biomass reactions and individually integrate them into the input model we are interested in. We propose the following steps:

1. Load, one by one, the models of interest on MATLAB. For example, if we focus on human GEM reconstructions, we could use Human 1 (Robinson et al., 2020), Recon 2 (Thiele et al., 2013), Recon 3D (Brunk et al., 2018), HMR 2.0 (Mardinoglu et al., 2014) or any other published model, based on our final objective. Depending on the type of file we have we can use either:

~~~
     > model = readCbModel(‘YOURMODEL.xml’,’fileType’,’SBML’);
~~~ Or

~~~
     > load YOURMODEL.mat
     > model = YOURMODEL;
~~~
2. Identify the biomass reaction defined in this model:

~~~
     > Biomass_idx = find(ismember(model.rxns,’NAME_BIOMASS_RXN’))
~~~ and visualise the formulation of the reaction of interest:

~~~
     > printRxnFormula(model, model.rxns(Biomass_idx))
~~~

**CRITICAL:** Remember to check in advance the name of the biomass reaction in the model under study, which needs to be inputted in step 2, NAME_BIOMASS_RXN.

### Step 2: Obtain a model-compatible biomass reaction

#### Timing: 15-30 min

To integrate a biomass reaction into a model, each metabolite in the biomass must be produced by reactions in the GEM. As GEMs often have different identifiers for metabolites and reactions, the identifier of the objective reaction had to be converted into the identifiers of the GEM. In our example, the identifiers from the Human 1 biomass are converted into Recon X identifiers. This standardisation step is done using Metabolic Atlas (https://metabolicatlas.org). For other GEMs and species, alternative databases might be more appropriate, such as the BiGG (http://bigg.ucsd.edu/), or VMH (https://www.vmh.life/). Alternatively, BiGG offers dictionaries for metabolites and reactions of several models that can be downloaded as a text file from their webpage: http://bigg.ucsd.edu/data_access. Despite the used resource, the principle is the same.

3. Select the Human repository in the Metabolic Atlas searching tool (Figure 2, red square).
4. Introduce one by one the identifiers of the metabolites in the biomass reaction under study in the Metabolic atlas search tool (Figure 2, blue square). For example, the human 1 biomass reaction is: *45* ***m01371c*** *+ 0*.*0267 m01721n + 45 m02040c + 0*.*1124 m02847c + 0*.*4062 m03161c + 0*.*0012 m10012c + 5*.*3375 m10013c + 0*.*2212 m10014c + 0*.*4835 m10015c --> 45 m02039c + 45 m02751c + temp001c*. Hence, we manually type the first identifier, m01371c, to see which is its equivalent in Recon 2 (Figure 3). Note that the names of the metabolites in the Metabolic Atlas site were updated between the time of the publication Moscardo Garcia *et al*., 2021 and the present paper. Metabolite’s identifiers of the type m followed by a number and the abbreviation of the compartment (i.e., m01371c) were changed to MAM followed by the previous identifier. Hence, the equivalent identifier for m01371c in the Metabolic Atlas database is MAM01371c. Then, the Recon X (or BIGG) compatible identifier is atp and, since we are talking about the cytosolic form, we would write atp[c]. In Recon2, the compartments are symbolised by their initial between square brackets i.e. [c] for cytosol, [n] for nucleus, [m] for mitochondria, [g] for Golgi, [e] for extracellular compartment, and [r] endoplasmic reticulum. In other models, other symbols are used like “_s” or “s” for the extracellular compartment in human1. Following this approach, the resulting human 1 biomass reaction is: *45 atp[c] + 0*.*0267 dna[c] + 45 h2o[c] + 0*.*1124 rna[c]* + *0*.*4062 glycogen[c] + 0*.*0012 cofactor_pool[c] + 5*.*3375 protein_pool[c] + 0*.*2212 lipid_pool[c] + 0*.*4835 metabolite_pool[c] --> 45 h[c] + 45 pi[c] + biomass[c]*. **Note:** Nevertheless, not all metabolites have a BiGG identifier, since they do not correspond to an actual metabolite but to a protein, RNA, or other more complex molecules. In these cases, the user can choose any suitable name without spaces or symbols. Later, the corresponding reactions producing these metabolites will be added. For example, in the Human 1 biomass reaction, many metabolites are not available as BiGG identifiers, such as *m02847c, m03161c, m10012c, m10013c, m10014c, m10015c* and *temp001c*. Hence, non-standard names were given to them, rna[c], glycogen[c], cofactor_pool[c], protein_pool[c], lipid_pool[c] and metabolite_pool[c], respectively.

**Figure 2.**
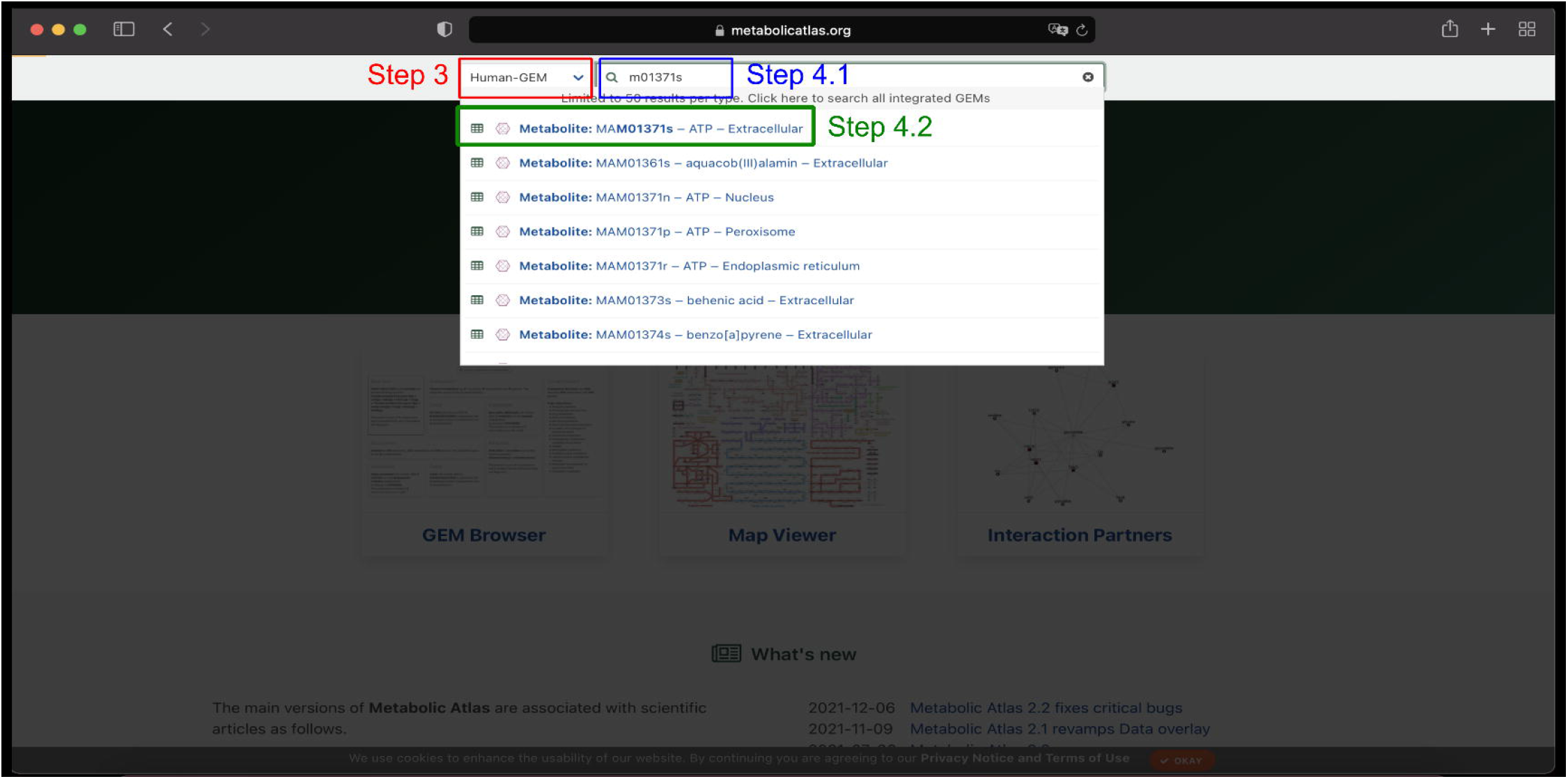
Select the Human-GEM repository. After accessing the Metabolic Atlas website (https://metabolicatlas.org), we need to select Human-GEM (red square) in the searching tool to start with the conversion of the identifiers. Afterwards, we can manually type the identifier we want to convert (blue square) and select it to go to its information page (green square).

**Figure 3.**
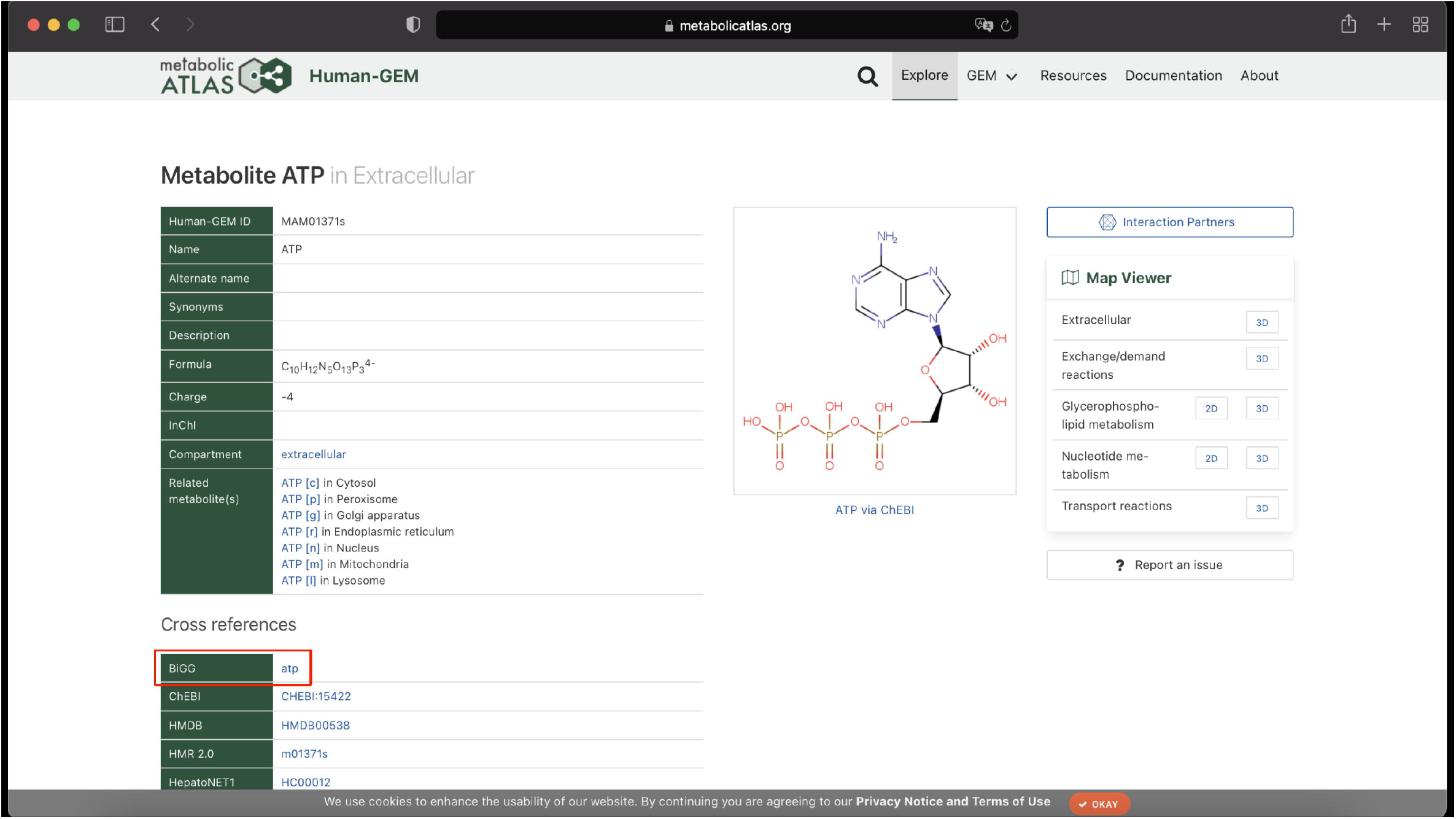
Build a Recon 2 compatible reaction. Each metabolite identifier is manually introduced into the metabolic atlas website. Then, by looking at the external databases information, a new reaction is built by considering the BiGG identifiers instead of those in the original reaction.

### Step 3: Model consistency

#### Timing: 1-2 hours

Once the reaction of interest has been formulated in a form compatible with the input model, we need to check if the model is consistent. In other words, if all reactions included in the model can carry a flux. If a metabolite that is included in the biomass formulation cannot be produced by the model due to gaps or dead-ends, the biomass will be blocked (unable to carry a flux) and the reaction will be removed by the consistency checking algorithm FastCC (Vlassis et al., 2014). By checking which of the biomass metabolites are not present in the model after the run of FastCC, allows pinpointing where the dead-ends are localised, notably in the production pathways of these metabolites.

5. First, remove the default biomass reaction (or any reaction that needs to be replaced) in the input model to avoid mixing reactions in the future and rename the new model, in this case model_no_Biomass:

~~~
> input_model = readCbModel(‘YOURINPUTMODEL.xml’,’fileType’,’SBML’);
> idx = find(∼cellfun(@isempty,regexpi(input_model.rxns,
‘NAMEREACTION’)));
> model_no_Biomass = removeRxns(input_model, input_model.rxns(idx));
~~~
6. Integrate the new reaction in the desired input model, model_no_Biomass:

~~~
> model_with_new_biomass = addReaction (model_no_Biomass,
‘biomass_human1’, ‘reactionFormula’,’0.0012 cofactor_pool[c] +
5.3375 protein_pool[c] + 0.1124 rna[c] + 0.0267 dna[c] + 0.2212
lipid_pool[c] + 0.4062 glycogen[c] + 0.4835 metabolite_pool[c] + 45
atp[c] + 45 h2o[c] -> biomass[c] + 45 pi[c] + 45 h[c]’);
~~~
7. Check the model consistency.

~~~
> A = fastcc_4_rfastcormics(model_with_new_biomass, 1e-4,0);
~~~ The output of this line of code is either a sentence “The input model is consistent” or a vector A with all the consistent reactions in the model. In the former case, since the model is already consistent, please, proceed directly to Step 4. Alternatively, in the latter case consistency needs to be achieved so, please, follow the next steps.
8. To achieve model consistency, we first need to check the metabolite included in the biomass formulation are present in the input model. To do so we can either check one by one the presence of each metabolite:

~~~
> find(ismember(model_no_Biomass.mets,’METABOLITENAME’))
~~~ or store the metabolites of the biomass function in a variable called Biomass_mets and compare it with the set of metabolites in our model:

~~~
> Biomass_mets = {‘METABOLITENAME1’,’METABOLITENAME2’};
> setdiff(Biomass_mets, model_no_Biomass.mets)
~~~
9. In case the metabolite is not found, a reaction producing such metabolite has to be added to achieve model consistency. In our example, the reactions are obtained from the Metabolic Atlas site (Figure 4). Once we search for the metabolite of interest, we focus on the reactions producing it. Introducing a new reaction can require the introduction of new metabolites and hence new reactions. Therefore, it is wise to prioritise reactions that included metabolites already present in the model or reactions that required the introduction of a few other reactions. Thus, to avoid an endless process, the user can check which threshold of additional reactions allows them to integrate most metabolites within a reasonable time, for example in this study a threshold of 4 additional reactions was heuristically defined for the present model and biomass. This number is dependent on the model and the users might need less or more reactions depending on the importance of the metabolite to be added. The presented protocol is an iterative process, and the users should consider the improvement of prediction power when adding or removing a metabolite. Hence, if a metabolite requires the addition of too many reactions, the metabolite is removed from the biomass formulation.

**Figure 4.**
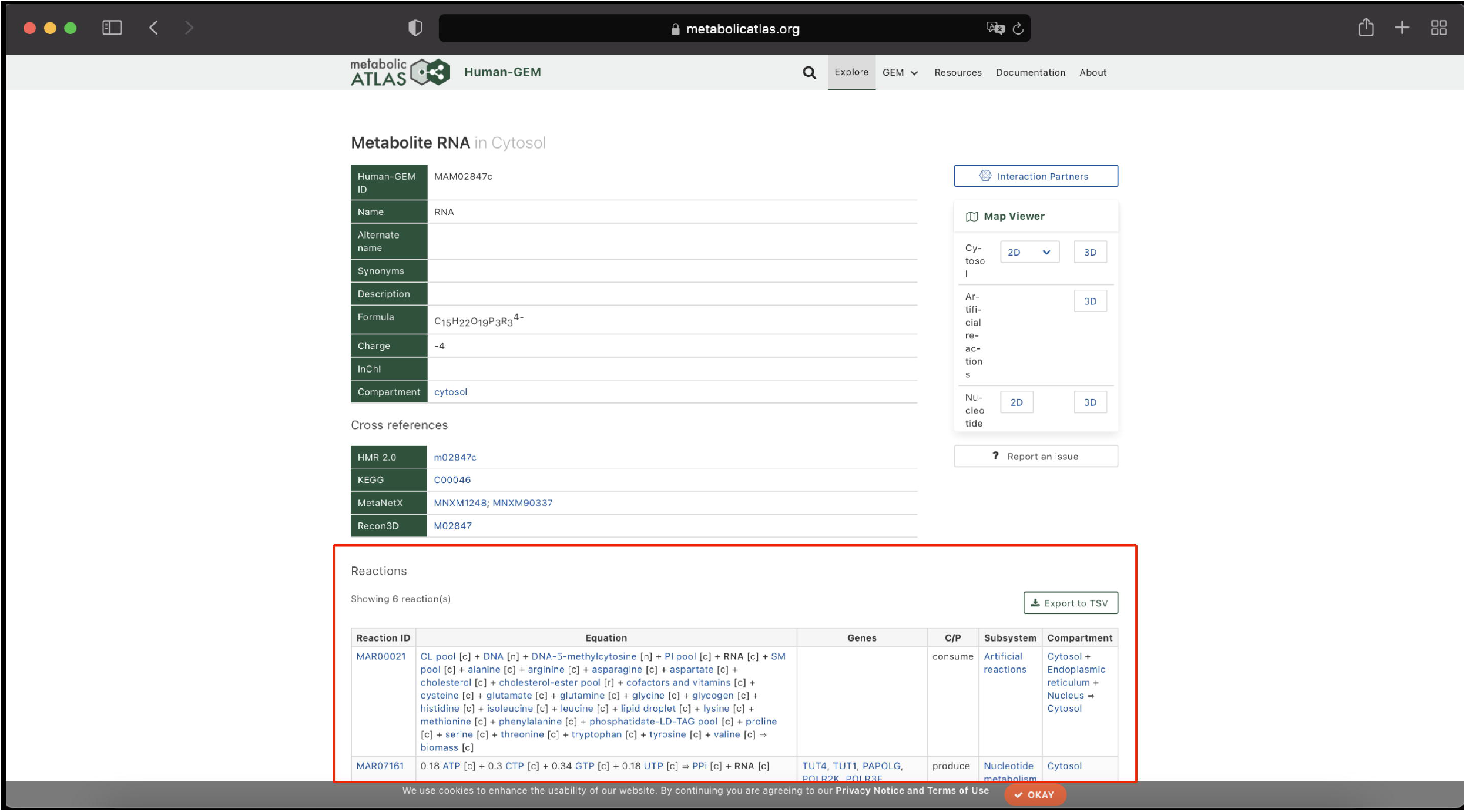
Introduction of the reaction producing the metabolite of interest to achieve model consistency. Once the metabolite site has been accessed, the Reactions section (red square) contains all the reactions producing or consuming the metabolite of interest. Thus, one of these reactions will be included in the model to overcome the inconsistency problem.

~~~
> model_with_new_biomass = addReaction (model_with_new_biomass,
‘REACTION_NAME’, ‘reactionFormula’,’0.0012 X[c] + 0.3375 Y[c] -> 0.5
MET_INTEREST[c]’);
~~~ Following the human 1 example, the metabolite *rna[c]* is not part of Recon 2. Thus, in the Metabolic atlas, we identify a reaction with 4 metabolites (all of them present in Recon 2) able to produce rna[c]. Hence, the reaction is included in the model as follows:

~~~
> model_with_new_biomass = addReaction (model_with_new_biomass,
‘RNA_production’,’reactionFormula’,’0.18 atp[c] +0.3 ctp[c] +0.34 gt
p[c] +0.18 utp[c] -> ppi[c] + rna[c]’);
~~~ This step is repeated for every metabolite in the reaction we want to integrate as well as for every metabolite from the new reactions we have included to achieve consistency. A script including the integration of all metabolites from Human 1 biomass reaction in Recon 2 can be found in https://github.com/sysbiolux/Biomass_formulation/tree/main/Scripts/Generation_Data_Human1_Figure1_4a.m. **Note**: From experience, we recommend including one metabolite in the biomass reaction and the required reactions for this specific metabolite at a time and checking the consistency after each metabolite addition. It is easier to identify the reactions leading to an inconsistent model.

#### Hint for advanced users

If a high number of dead ends and gaps prevent the inclusion of the reaction of interest, a gap-filling algorithm might be considered, such as fastGapfill (Thiele et al., 2014). However, to avoid the inclusion of reactions with low confidence, whose addition would decrease the prediction power of the network, it is crucial to manually curate the network after this process. In networks that are highly curated and require a low number of metabolites to be included, a manual inclusion should be preferred over an automated one.

When setting up a biomass reaction, the procedure described in Chan et al., 2017 should be followed to achieve a standardisation of the growth to a molecular weight (MW) of 1□g mmol^−1^. The authors give a respective MATLAB code which can be combined with our pipeline. Importantly, for the purpose of our pipeline which is to transfer an existing biomass function to another model, if metabolites are removed or added to the biomass function, the coefficients need to be rescaled to maintain the given MW of the biomass function. Assuming that the original MW is 1□g mmol^−1^ (i.e., the Chan et al procedure was executed), the addition of metabolites i and removal of metabolites j will change the growth MW to:

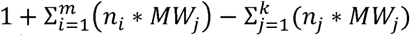

Rescaling to 1□g mmol^−1^ is obtained by dividing all coefficients of the biomass function (not of the biomass metabolite itself, if existing) by this factor. To be consistent the MW of the metabolites must be given in g mmol^−1^. Hence, the biomass reaction with the new coefficients will be added (c1-c10 stand for the new calculated values).

~~~
> model_with_new_biomass = removeRxns (model_with_new_biomass, NAME_BIOMASS_RXN).
> model_with_new_biomass = addReaction (model_no_Biomass,
‘biomass_human1’, ‘reactionFormula’,’c1 cofactor_pool[c] + c2
protein_pool[c] + c3 rna[c] + c4 dna[c] + 0.2212 lipid_pool[c] + c5
glycogen[c] + c6 metabolite_pool[c] + c7 atp[c] + c8 h2o[c] ->
biomass[c] + c9 pi[c] + c10 h[c]’);
> model_with_new_biomass.rev(end)=0;
~~~

10. After including the biomass reaction, or any reaction, of interest and all the reactions required for consistency, check the model consistency again via fastcc_4_rfastcormics.

~~~
> A = fastcc_4_rfastcormics(model_with_new_biomass, 1e-4,0);
~~~ At this point, the output should be “The input model is consistent”, meaning that we can proceed to test the performance of the model with the new set of reactions. The consistent model obtained through these steps has been deposited in https://github.com/sysbiolux/Biomass_formulation/tree/main/MethodsPaper/H1Bioma_ss_R2.mat.

### Step 4: Test model performance

#### Timing: 30 min

After obtaining the desired consistent input model with the new reaction of interest, we can test its predictive capacity. Although there are different ways to evaluate that, our approach is to determine the sensitivity, specificity, and precision of the essential gene predictions against CRISPR high-throughput data. Additionally, we explore the predictive capacity of the new model in terms of growth rate predictions.

11. First of all, the gene expression data of interest is loaded. As previously mentioned, this data should be provided in matrix form, in which the rows and columns correspond to the genes and samples respectively. The names of the samples correspond to cell lines, culture conditions or replicates and are provided as a cell array called colnames. While rownames correspond to a cell array that includes the gene names.

~~~
> load DATAOFINTEREST.mat
~~~ The data should be imported into matlab and split into 3 variables: fpkm, colnames, and rownames.

- colnames is cell array containing the names of the samples;
- fpkm is a matrix containing as many lines as the rownames and as many columns as the samples/replicates.
- rownames is a cell array that contains the same number of lines and contains the data identifier i.e., gene symbols or Ensembl ID or Entez id;

~~~
> %Discretise the data
> discretized = discretize_FPKM(fpkm, colnames);
~~~ **Note:** a discretisation is needed as rFASTCORMICS only recognises two states for the reactions in the model, present and absent, and 3 states for the genes in the gene expression dataset, expressed, not expressed and unknown. The discretisation step allows determining for each sample the expression threshold above which a gene is considered to be expressed and the threshold below which a gene is not expressed. Genes with values between the 2 thresholds are considered undermined. The gene expression status will be mapped to the reactions to obtain a set of inactive reactions that will be included in the output model, and core reactions that will be included.
12. The users need to produce a dictionary, a table with 2 columns that maps the model gene identifiers and the data identifiers. The first column should match the model identifier whereas the second column contains the data identifier. Online tools such as biodbNET (Mudunuri et al., 2009) or R tools like bioMART (Durinck et al., 2009) can be useful to create these dictionaries. > load DICTIONARY.mat
13. Context-specific models, which contains reactions active in the context of interest, are obtained by integrating transcriptomic data (i.e., CCLE data Barretina et al., 2019) into the new model we obtained in previous steps using rFASTCORMICS (Pacheco et al., 2019). In this case, since we want to assess the performance of the new biomass reaction as objective function, we define as objective function the reaction we have described in Step 6, biomass_human1. The optional_settings is an object that can include several fields such as medium, func and not_ medium_constrained. Usually, we recommend the user to include the medium composition that defines the set of metabolite that can be uptake by the model (optional_settings.medium) as well as the non-constraint reactions, if needed, metabolites that are not found in the media composition used experimentally but are required due to model shortcoming to allow the objective to carry a flux (optional_settings.not_medium_constrained) and the objective function of interest (optional_settings.func). rFASTCORMICS allows building consensus and sample specific models. For the building of consensus models where data for several samples or patients are pooled, a consensus proportion is set that defines the fraction of the pooled samples (recommended value 0.9) that must express or not to express a gene in order to call it as expressed or not expressed in the model. The option already_mapped is set to 1 if the user has already mapped the expression data to the reactions of the model reactions otherwise rFASTCORMICS will perform the mapping using the Gene Protein Rules (GPR) of the model. Epsilon is the threshold below which a flux will be considered to be equal to zero.

~~~
> optional_settings.func = {‘biomass_human1’};
> biomass_reaction = {‘biomass_human1’};
> epsilon = 1e-4;
> consensus_proportion = 0.9;
> already_mapped_tag = 0;
> [∼, A] = fastcormics_RNAseq(model_with_new_biomass, discretized,
rownames, DICTIONARY,biomass_reaction, already_mapped_tag,
consensus_proportion, epsilon, optional_settings); %if this line
lead to the error: “Error: The input was too complicated or too big
for MATLAB to parse”. Please run line 5
~~~ **Note**: The objective function can be adjusted to the interest of our study. In this case, the name of the reaction will be changed accordingly:

~~~
> optional_settings.func = {‘NAME_OBJECTIVE_FUNCTION’};
~~~ Constraints, such as medium composition, can be considered for the model reconstruction by defining the available metabolites in the medium. The metabolite identifiers present in the medium are given as a list and stored in optional_setting.medium. See Moscardo Garcia et al., (2021) for more details.

~~~
> optional_settings.medium = {‘ID_METABOLITE_IN_MEDIA1; ID_METABOLITE_IN_MEDIA2’};
~~~ **Note:** The ‘ID_METABOLITE_IN_MEDIA1’ should match the identifier model.mets.
14. Perform single gene deletion to obtain a list of predicted essential genes.

~~~
> model_out = removeRxns(model_with_new_biomass,
model_with_new_biomass.rxns(setdiff(1:numel(model.rxns), A)));
>  model_out = changeObjective(model_out,NAME_OBJECTIVE_FUNCTION);
> [grRatio, grRateKO, grRateWT, hasEffect, delRxns, fluxSolution,
genelist] = singleGeneDeletion(model_out,’FBA’,[],0,1);*
> Predicted_Essential(1).geneList=genelist(find(grRatio<THRESHOLD));
~~~ * As the inputs and outputs of the *singleGeneDeletion* function of the COBRA might change, we recommend the version provided with our code. The THRESHOLD is a number between 0 and 1, which is user defined and corresponds to the ratio below which the knock-out of a gene is considered essential.
15. Experimental data is required for validation. CRISPR-Cas 9 data can be used to validate the set of predicted essential genes whereas experimentally measured growth rates are required for the validation of the growth rate predictions. The CRISPR-Cas 9 data is expected in a binary table form with genes in the rows and samples in the columns, where 0 stands for not expressed and 1 for expressed. Alternatively, growth rates should be provided in a table form with rows representing the analysed samples and one column with experimental doubling times. First, compare the predicted essential genes with experimentally identified essential genes, to determine the precision, sensitivity and specificity of the predictions. In the following script a gene is considered essential if a knock-out reduces the biomass production by half (threshold 0.5).

~~~
>  confusion_table=computeEnrichment_tests(Predicted_Essential,
colnames, data_essential, colnames_essential, rownames_essential,
dico_essential, genelist);
confusion_table.Specificity
confusion_table.Precision
confusion_table.Sensitivity
~~~ DATA_ESSENTIAL is a binary matrix; the columns correspond to samples and the row to genes. 1s in the matrix correspond to essential genes and 0s to not essential. COLUMNNAMES_ESSENTIAL is a cell array with the sample’s names and ROWNAMES_ESSENTIAL with the gene identifiers. DICO_ESSENTIAL matches the identifiers used in the essential gene datasets and the model.
16. Additionally, the predictive capacity of the model can also be assessed in terms of growth rate predictions.

~~~
> ind = find(∼cellfun(@isempty,regexp(model_with_new_biomass.rxns,
optional_settings.func{1})));
> model_out = removeRxns(model_with_new_biomass,
model_with_new_biomass.rxns(setdiff(1:numel(model_with_new_biomass.rx
ns),A)));
> model_out = changeObjective(model_out,
model_with_new_biomass.rxns(ind));
> sol = optimizeCbModel(model_out);
> optimization_results = sol.f;
~~~ These numerical values are directly compared with experimentally measured growth rates to determine how far the predictions are from the experimental values. **Note:** Additional constraints regarding exometabolomic data can be included to simulate the experimental conditions under study. See Moscardo Garcia et al., (2021) for more details.

### Expected outcomes

In this protocol, a new consistent model containing reactions coming from other GEMs is obtained. In this particular case, the new reaction was defined as the objective function. Thus, we provide two possibilities to assess the performance of the new model: essential gene and growth rate predictions.

### Limitations

#### Lack of context-specific biomass formulations

At the present moment, several biomass functions were published but most of them are generic functions. Hence, they are not specific to a given cell type or a given context such as high proliferative cells and cancer cells. Even for the generic models, biomasses are often adapted from other organisms, or a draft biomass is automatically generated for which the coefficients were adapted to the organism.

However, a cancer cell that relies on glycolysis should not have the same biomass formulation as a healthy cell or a cancer cell that rely on oxidative phosphorylation. Several papers especially working on yeast or *E. coli* discuss how biomass composition and coefficient affect flux distributions. Hence, several initiatives were launched to measure experimental fluxes in these two organisms. However, similar attempts, even if necessary, might be more difficult in cancer, as multicellular organisms are more complex. The interplay between the cancer cells, the surrounding tissues, and the microenvironment, would need to be considered in the growth optimisation, as some biomass metabolites might be produced by cancer-associated fibroblast or other surrounding cells. Further, it was shown that biomass formulation in yeast and E. coli should be tailored for different strains. The same is true for different cancer subtypes and potentially for the clones within a tumour.

Nevertheless, there is space for improvement and need for integration methods for different types of data such as lipidomic among others. Recent efforts have been made to include the diversity of lipids inside GEMs but also in the biomass formulation, such as SLIME (Sánchez et al., 2019) to integrate different classes of lipids via the use of SLIME reactions that split reactions in lipids component but also consider acyl distributions to impose fluxes. Other metabolite types could be included in the different biomass formulations depending on the needs of the studied organism. Hence to decide on the biomass composition and coefficients based on experimental data, BOFdat (Lachance et al., 2019), a tool to semi-automatedly reconstruct a biomass function using, among others, several types of omics data and a generic algorithm was published. However, the tool requires lots of different data types that are not always available for all studied contexts (Moscardo Garcia et al., 2021).

Another issue is the lack of standardization in GEM in general but specifically also of the biomass formulation. Reactions should be balanced in mass and in charge. But, although it is often stated in papers that reactions are mass and charged balanced, it is not always the case. Unbalanced reactions often result from the inclusion of “artificial” metabolites required for modelling purposes. These do not represent an actual metabolite, but a more complex entity composed of various metabolites such as protein, RNA, tRNA-alanine, etc. Hence, they do not have formulas (or having X or Y in the metFormulas field). Tools such as Memote (Lieven et al, 2020) that allow assessing the quality of metabolic models, among others, in terms of annotation and consistency, contribute to the obtention of better biomass formulations. Also, efforts to standardise identifiers such as MetaNetX (Moretti et al, 2021, www.metanetx.org) that allow comparing models among them and to external databases, facilitate the spotting of errors in metabolic models. However, more standardisation efforts are needed to allow i.e., mapping the biomass metabolites without having to search external databases. Ideally, a field including an unambiguous identifier, that would allow mapping metabolites across models of different organisms should be mandatorily included in every model. Furthermore, guidelines on the use of metabolites with side groups should be established to allow comparing the different biomass formulations. Finally, the biomasses should be standardised to produce 1g of biomass per mmol of metabolites. This is particularly important in the modelling of bacterial communities, for which non-standardised biomasses could bias the ratio of the modelled species.

#### Lack of automatization of optimization of the biomass formulation

The workflow only describes how to add a biomass function but does not show how to create or tailor a biomass function to the context of interest. Since alternative pathways producing some components of the biomass might be missing in the genome-scale reconstruction, a knock-out of the genes implicated in the producing pathways might reduce the growth *in silico*. However, *in vivo* the knockouts would have been compensated by alternative pathways not present in the model, causing an increase in the number of False Positives. Thus, in some cases, removing the metabolite from the list of biomass components would result in better accuracy.

#### Gene Protein Reaction Rules

The Gene Protein Reaction (GPR) rules code the relation between reaction and genes, meaning which genes are participating in the regulation of each reaction. Several genes can control the same reaction. If the genes are isozymes, only one gene is required to activate the reaction and will be encoded in the GPR rules with a BOOLEAN OR (for instance, A OR B means that the reaction is controlled by the isozymes A and B, and only one must be expressed to have an active reaction). If the genes form a complex, A BOOLEAN AND will be used and both genes must be expressed for the reaction to be active. However, differences in terms of the GPR rules can be found in currently published models. Hence, the lack of consistency in the GPR rules definition can lead to significant differences in the results.

#### Scarcity of experimental data for some organism

The workflow requires high-throughput screen data or experimentally identified essential genes in the context of interest to validate the predicted essential genes as well as flux rate data with the upper and lower bounds. In some cases, this type of data is not available for the context of interest which may restrict the validation of our results.

#### Troubleshooting

##### <u>Problem</u> 1

The input model is missing grRules, which allow us to determine the reactions present after the model reconstruction.

###### Potential solution

This problem can be solved using the function generateRules_rFASTCORMICS.m, which can be retrieved from https://github.com/sysbiolux/rFASTCORMICS

##### Problem 2

In some systems step 13 can lead to the error: “*Error: The input was too complicated or too big for MATLAB to parse”.*

###### Potential solution

In this case, please add

~~~
> feature astheightlimit 2000
~~~

at the beginning of your code.

##### Problem 3

I am interested in using a context-specific algorithm other than rFASTCORMICS.

###### Potential solution

Recommended for experts in context-specific reconstructions only. Run the algorithm you are interested in with the required inputs instead of Step 13, and run Steps 14 to 16 on the output model generated by the context-specific building algorithm. Numerous implementations are available in supplements of benchmarking papers and in the Cobra toolbox. Note that some implementations are different from the original papers, which might affect the quality of the results. The quality of the context-specific reconstruction is strongly affected by the choice of the algorithm. Some algorithms do not provide a discretisation step, however it is a critical step for most algorithms and arbitrary thresholds should never be used, as the quality of the context-specific reconstruction is strongly affected by the used threshold.

##### Problem 4

I would like to use a solver alternative to Cplex.

###### Potential solution

Cplex is free for academics. However, other solvers have a version compatible with the Cobra toolbox which can be downloaded from GitHub https://github.com/sysbiolux/rFASTCORMICS.

##### Problem 5

Pools of metabolites present in the biomass reaction have side groups denoted as R in the metabolite formula, which restrict the obtention of the molecular weights for the standardisation of the biomass reaction.

###### Potential solution

In this case, the metabolite formula should be replaced by the full side chain instead of R. The metabolite formula can be obtained from any database such as KEGG and Metabolic Atlas among others. Alternatively, we refer the author to Chan et al., 2017 (https://github.com/maranasgroup/BiomassMW/tree/master/MatlabCobraToolbox), who provided MATLAB scripts to obtain the metabolite formula and the corresponding MW.

## Resource availability

### Lead contact

Further information and requests for resources and reagents should be directed to and will be fulfilled by the lead contact, Professor Thomas Sauter.

### Materials availability

This study did not generate new unique reagents.

### Data and code availability

- This paper analyses existing, publicly available data. The source of the data is listed in the key resources table.
- A standard template to apply this protocol can be found in our GitHub repository as Template_code.m (https://github.com/sysbiolux/Biomass_formulation/tree/main/MethodsPaper/) and is publicly available as of the date of publication.
- Any additional information to reproduce the results reported in this paper is available from the lead contact upon request.

## Acknowledgments

This study was supported by the University of Luxembourg and the Luxembourg National Research Fund (FNR PRIDE PRIDE15/10675146/CANBIO).

## Author contributions

M.M.G carried out the experiments. M.P.P and T.S conceived and planned the experiment.

M.M.G and M.P.P wrote the manuscript. All the authors read and revised the manuscript.

## References

Agren, R., Bordel, S., Mardinoglu, A., Pornputtapong, N., Nookaew, I., & Nielsen, J. (2012). Reconstruction of Genome-Scale Active Metabolic Networks for 69 Human Cell Types and 16 Cancer Types Using INIT. In C. D. Maranas (Ed.), PLoS Computational Biology (Vol. 8, Issue 5, p. e1002518). Public Library of Science (PLoS). https://doi.org/10.1371/journal.pcbi.1002518.

Barretina, J., Caponigro, G., Stransky, N., Venkatesan, K., Margolin, A. A., Kim, S., Wilson, C. J., Lehár, J., Kryukov, G. V., Sonkin, D., Reddy, A., Liu, M., Murray, L., Berger, M. F., Monahan, J. E., Morais, P., Meltzer, J., Korejwa, A., Jané-Valbuena, J., Mapa, F. A., … Garraway, L. A. (2012). The Cancer Cell Line Encyclopedia enables predictive modelling of anticancer drug sensitivity. Nature, 483(7391), 603–607. https://doi.org/10.1038/nature11003.

Brunk, E., Sahoo, S., Zielinski, D. C., Altunkaya, A., Dräger, A., Mih, N., Gatto, F., Nilsson, A., Preciat Gonzalez, G. A., Aurich, M. K., Prlić, A., Sastry, A., Danielsdottir, A. D., Heinken, A., Noronha, A., Rose, P. W., Burley, S. K., Fleming, R. M. T., Nielsen, J., … Palsson, B. O. (2018). Recon3 enables a three-dimensional view of gene variation in human metabolism. Nature Biotechnology, 36(3), 272–281. https://doi.org/10.1038/nbt.4072

Chan, S. H. J., Cai, J., Wang, L., Simons-Senftle, M. N., & Maranas, C. D. (2017). Standardizing biomass reactions and ensuring complete mass balance in genome-scale metabolic models. In J. Wren (Ed.), Bioinformatics (Vol. 33, Issue 22, pp. 3603–3609). Oxford University Press (OUP). https://doi.org/10.1093/bioinformatics/btx453

DepMap Achilles 19Q1. (2019). Public. https://doi.org/10.6084/m9.figshare.7655150.v1.

Durinck S., Spellman P., Birney E., Huber W. (2009). Mapping identifiers for the integration of genomic datasets with the R/Bioconductor package biomaRt. Nature Protocols, 4, 1184–1191.

Jain, M., Nilsson, R., Sharma, S., Madhusudhan, N., Kitami, T., Souza, A. L., Kafri, R., Kirschner, M. W., Clish, C. B., & Mootha, V. K. (2012). Metabolite Profiling Identifies a Key Role for Glycine in Rapid Cancer Cell Proliferation. Science, 336(6084), 1040–1044. https://doi.org/10.1126/science.1218595

Lachance, J.-C., Lloyd, C. J., Monk, J. M., Yang, L., Sastry, A. v., Seif, Y., Palsson, B. O., Rodrigue, S., Feist, A. M., King, Z. A., & Jacques, P.-É. (2019). BOFdat: Generating biomass objective functions for genome-scale metabolic models from experimental data. PLOS Computational Biology, 15(4), e1006971. https://doi.org/10.1371/journal.pcbi.1006971

Lieven, C., Beber, M.E., Olivier, B.G. et al. MEMOTE for standardized genome-scale metabolic model testing. Nat Biotechnol 38, 272–276 (2020). https://doi.org/10.1038/s41587-020-0446-y

Mardinoglu, A., Agren, R., Kampf, C., Asplund, A., Uhlen, M., Nielsen, J. (2014). Genome-scale metabolic modelling of hepatocytes reveals serine deficiency in patients with non-alcoholic fatty liver disease. Nature Communications, 5(1), 3083. https://doi.org/10.1038/ncomms4083

Moretti, S., Tran, V. D. T., Mehl, F., Ibberson, M., & Pagni, M. (2020). MetaNetX/MNXref: unified namespace for metabolites and biochemical reactions in the context of metabolic models. In Nucleic Acids Research (Vol. 49, Issue D1, pp. D570–D574). Oxford University Press (OUP). https://doi.org/10.1093/nar/gkaa992

Moscardó García M., Pacheco M., Bintener T., Presta L., Sauter T. (2021). Importance of the biomass formulation for cancer metabolic modeling and drug prediction. iScience. 10;24(10):103110. https://doi.org/10.1016/j.isci.2021.103110

Mudunuri, U., Che, A., Yi, M. and Stephens, R.M. (2009) bioDBnet: the biological database network. Bioinformatics, 25, 555–556.

Pacheco, M. P., Bintener, T., Ternes, D., Kulms, D., Haan, S., Letellier, E., Sauter, T. (2019). Identifying and targeting cancer-specific metabolism with network-based drug target prediction. EBioMedicine, 43, 98–106. https://doi.org/10.1016/j.ebiom.2019.04.046

Pacheco, M.P., John, E., Kaoma, T., …, Sauter, T. (2015). Integrated metabolic modelling reveals cell-type specific epigenetic control points of the macrophage metabolic network. BMC Genomics, 16, 809 https://doi.org/10.1186/s12864-015-1984-4

O’Connor, R., Kauffmann-Zeh, A., Liu, Y., Lehar, S., Evan, G., Baserga, R., Blättler, R. (1997). The IGF-I receptor domains for protection from apoptosis are distinct from those required for proliferation and transformation. Mol. Cell. Bio, 17, 427–435.

Robinson, J. L., Kocabaş, P., Wang, H., Cholley, P.-E., Cook, D., Nilsson, A., Anton, M., Ferreira, R., Domenzain, I., Billa, V., Limeta, A., Hedin, A., Gustafsson, J., Kerkhoven, E. J., Svensson, L. T., Palsson, B. O., Mardinoglu, A., Hansson, L., Uhlén, M., Nielsen, J. (2020). An atlas of human metabolism. Science Signaling, 13(624), eaaz1482. https://doi.org/10.1126/scisignal.aaz1482

Sánchez, B.J., Li, F., Kerkhoven, E.J. et al. SLIMEr: probing flexibility of lipid metabolism in yeast with an improved constraint-based modeling framework. BMC Syst Biol 13, 4 (2019). https://doi.org/10.1186/s12918-018-0673-8

Thiele I., Swainston N., Fleming R. M., Hoppe A., Sahoo S., Aurich M. K., Haraldsdottir H., Mo M. L., Rolfsson O., Stobbe M. D., Thorleifsson S. G., Agren R., Bölling C., Bordel S., Chavali A. K., Dobson P., Dunn W. B., Endler L., Hala D., Hucka M., Hull D., Jameson D., Jamshidi N., Jonsson J. J., Juty N., Keating S., Nookaew I., Le Novère N., Malys N., Mazein A., Papin J. A., Price N. D., Selkov E., Sigurdsson M. I., Simeonidis E., Sonnenschein N., Smallbone K., Sorokin A., van Beek J. H., Weichart D., Goryanin I., Nielsen J., Westerhoff H. V., Kell D. B., Mendes P., Palsson B. Ø. (2013) A community-driven global reconstruction of human metabolism. Nature Biotechnology. 31:419–425.

Thiele, I., Vlassis, N., & Fleming, R. M. (2014). fastGapFill: efficient gap filling in metabolic networks. Bioinformatics (Oxford, England), 30(17), 2529–2531. https://doi.org/10.1093/bioinformatics/btu321.

Wang, Y., Eddy, J.A. & Price, N.D. Reconstruction of genome-scale metabolic models for 126 human tissues using mCADRE. BMC Syst Biol 6, 153 (2012). https://doi.org/10.1186/1752-0509-6-153.

Vlassis, N., Pacheco, M. P., Sauter, T. (2014) Fast Reconstruction of Compact Context-Specific Metabolic Network Models. PLoS Comput Biol, 10(1): e1003424. https://doi.org/10.1371/journal.pcbi.1003424

